# Iron content of commercial mucin contributes to compositional stability of a cystic fibrosis airway synthetic microbiota community

**DOI:** 10.1101/2024.09.06.611695

**Authors:** Emily Giedraitis, Rachel L. Neve, Vanessa V. Phelan

## Abstract

*In vitro* culture models of mucosal environments are used to elucidate the mechanistic roles of the microbiota in human health. These models often include commercial mucins to reflect the *in-situ* role of mucins as an attachment site and nutrient source for the microbiota. Two types of mucins are commercially available: porcine gastric mucin (PGM) and bovine submaxillary mucin (BSM). These commercial mucins have been shown to contain iron, an essential element required by the microbiota as a co-factor for a variety of metabolic functions. In these mucin preparations, the concentration of available iron can exceed physiological concentrations present in the native environment. This unexpected source of iron influences experimental outcomes, including shaping the interactions between co-existing microbes in synthetic microbial communities used to elucidate the multispecies interactions within native microbiota. In this work, we leveraged the well-characterized iron-dependent production of secondary metabolites by the opportunistic pathogen *Pseudomonas aeruginosa* to aid in the development of a simple, low-cost, reproducible workflow to remove iron from commercial mucins. Using the mucosal environment of the cystic fibrosis (CF) airway as a model system, we show that *P. aeruginosa* is canonically responsive to iron concentration in the chemically defined synthetic CF medium complemented with semi-purified PGM, and community composition of a clinically relevant, synthetic CF airway microbial community is modulated, in part, by iron concentration in PGM.

**IMPORTANCE:** Mucins are critical components of *in vitro* systems used to model mucosal microbiota. However, crude commercial mucin preparations contain high concentrations of iron, which impacts interactions between members of the microbiota and influences interpretation of experimental results. Therefore, we developed and applied a simple, reproducible method to semi-purify commercial porcine gastric mucin as an affordable, low-iron mucin source. The development of this simplified workflow for semi-purification of commercial mucin enables researchers to remove confounding iron from a critical nutrient source when modeling clinically relevant microbial communities *in vitro*.

## INTRODUCTION

The mucosa is the largest barrier in the human body, lining the respiratory, digestive, and urogenital tracts. Contributing to this barrier is a mucus layer that covers the apical surface, preventing direct interactions of pathogenic microorganisms with the epithelia and providing an ecological niche for some members of the microbiota.(1, 2) The glycans of the large, heavily glycosylated mucins, the primary component of mucus, contribute to the composition and function of the microbiota.(3, 4) A proportion of the microbiota encode enzymes that degrade mucin, supporting a stable microbial community through metabolic cross-feeding.(5–8) To model the interactions between members of the mucosal microbiota *in vitro*, commercial mucins are often used.(9–12) Commercial mucins are the most economic choice as isolation of native mucins can be cost prohibitive and of limited quantity. Despite reduced viscoelastic, lubrication, and antimicrobial properties compared to their native counterparts, commercial mucins are useful for modeling metabolic cross-feeding within model synthetic microbial communities (SynComs).(13–16)

Two commercial mucins are used to model mucosal environments *in vitro*: bovine submaxillary mucin (BSM) and porcine gastric mucin (PGM).(9–12) Despite being used interchangeably, the mucin structures of BSM and PGM are different; BSM is composed of MUC5B and PGM is predominantly composed of MUC5AC.(17–21) Crude PGM and BSM also contain non-mucin components, including proteins, lipids, cellular debris, nucleotides, and iron. (11, 22–24) The undefined iron in commercial mucin likely influences the outcomes of microbial interactions under investigation in *in vitro* model systems. Iron is an essential nutrient required for microbial DNA replication, respiration, enzyme function, and growth.(25) In the human body, most iron is bound, leaving unbound iron concentrations below those required for metabolism and replication of bacterial cells.(26) This nutritional immunity induces pathogens to produce extracellular siderophores to bind ferric iron.(27) These siderophores capture iron from host iron-binding proteins and limit iron acquisition of co-occurring microbes, resulting in reduced viability of competitors.(28) We recently demonstrated that the concentration of iron in PGM is sufficient to support the growth of the opportunistic pathogen *Pseudomonas aeruginosa*, but significantly reduces its siderophore biosynthesis.(11)

The aims of this work were to determine the differential effect of BSM and PGM on the production of secondary metabolites by *P. aeruginosa* and establish the influence of mucin-associated iron on a four-member model CF airway synthetic microbial community (CF SynCom).(29) We chose *P. aeruginosa* and the CF SynCom as a model system for this study because the CF airway is colonized by a diverse microbiota resulting from defective mucociliary clearance and the structures, regulation, and functions of *P. aeruginosa* small molecule virulence factors are well characterized, including changes in metabolite abundance due to variations in iron availability.(30–32) Using untargeted metabolomics, we show that *P. aeruginosa* secondary metabolite production in synthetic CF medium 2 (SCFM2) is heavily impacted by the iron in commercial mucins, but mucin structure also plays a role in regulating the production of the redox active phenazines.(33) We demonstrate that the iron in commercial mucins can be easily removed by ultracentrifugation and dialysis and that the iron concentration in crude PGM contributes to stability of the CF SynCom in SCFM2 by preventing iron sequestration from *Staphylococcus aureus* by *P. aeruginosa*.

## RESULTS AND DISCUSSION

### Commercial mucins influence *P. aeruginosa* secondary metabolite production in an iron-dependent manner

As the research community moves towards elucidating the functions of the microbiota in health and disease, it is critical to evaluate the *in vitro* systems used to test hypotheses generated from *in vivo* data, including the media used for cultivation. Although chemically defined media is optimal for fine control of the nutrient environment, the complexity of the microbiota’s native mucosal environment and the limited quantity of pure mucin isolated from native sources to model *in vivo* conditions complicates the ability to implement ‘perfect’ *in vitro* systems. Due to the reliance of researchers on commercial materials to model mucosal environments *in vitro*, it was necessary to evaluate the effect of two common commercial mucins, BSM and PGM, on microbial physiology.

BSM and PGM are composed of different mucins and the commercial mucins have been shown to contain markedly different concentrations of iron.(11, 17–21, 24) Therefore, we sought to determine whether the type of crude mucin included in SCFM2 influenced the growth of P. aeruginosa PAO1 and its production of secondary metabolites. We chose *P. aeruginosa* as our model organism for this investigation for several reasons. It is an opportunistic pathogen that can extensively colonize the mucosal surfaces of the airways of people with CF, the impact of iron concentration on the regulation of its secondary metabolite biosynthetic pathways is well described, and analytical methods exist for capturing its small molecule chemical diversity.(11, 31, 34) The secondary metabolome of *P. aeruginosa* consists of a small number of molecular families with characterized roles in virulence, including functioning as mediators of quorum sensing, electron cycling, microbial competition, and iron acquisition.(30) Simultaneous measurement of these metabolites using untargeted metabolomics has been applied to identify biomarkers of virulence, genomic traits of clinical isolates, differential response to nutrient conditions, effects of disruption of biosynthetic pathways, and *in vivo* virulence factor production. (11, 35–38) We postulated that secondary metabolite profiling of PAO1 in otherwise chemically defined media would provide insight into whether the differential iron content of BSM and PGM is the primary driver of differences in levels of its small molecule virulence factors.

To test this hypothesis, we cultured PAO1 in SCFM2 without mucin (None-SCFM2) or complemented with either BSM or PGM. SCFM2 is a predominantly chemically defined medium comprised of amino acids and salts at concentrations measured from CF airway samples.(33) It is complemented with two chemically undefined components: DNA and mucin. In the original formulation of SCFM2, BSM was used as the mucin source.(33) However, PGM is commonly substituted for BSM, likely due to reduced cost and frequency of use in other CF artificial sputum media (ASM) formulations.(11, 29) Enumeration of viable cells after 48 hr of static incubation revealed that recovery of PAO1 from PGM-SCFM2 cultures was ∼3-fold higher than cultures in None- and BSM-SCFM2, suggesting that the nutrients in PGM enabled a slight growth advantage (Figure 1A).

**Figure 1.**
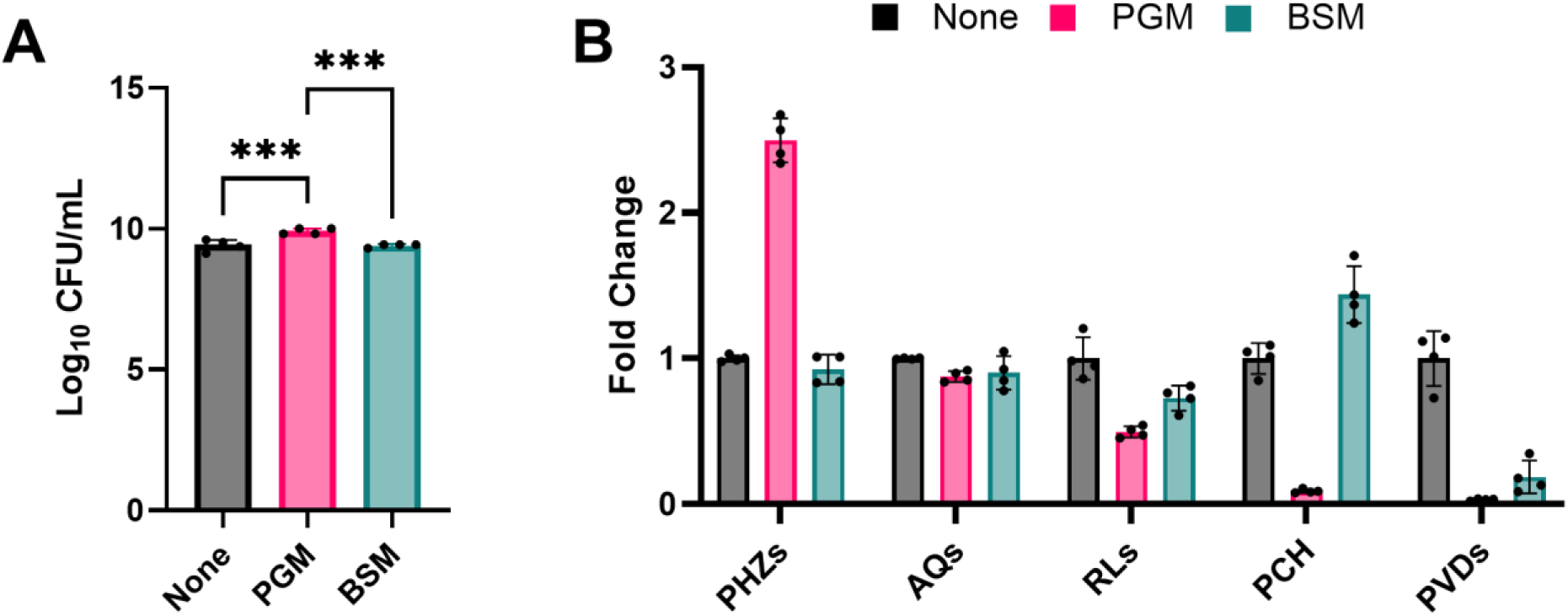
(A) Viable cell counts (CFU/mL) from *P. aeruginosa* PAO1 48 hr static cultures in None-SCFM2, PGM-SCFM2, and BSM-SCFM2. PGM-SCFM2 and BSM-SCFM2 contain 5 mg/mL crude mucin. (B) Fold change secondary metabolite levels produced by PAO1 in PGM- and BSM-SCFM2 compared to None-SCFM2. PGM: porcine gastric mucin; BSM: bovine submaxiliary mucin; PHZs: phenazines; AQs: alkyl quinolones; RLs: rhamnolipids; PCH: pyochelin; PVDs: pyoverdines. Corresponding abundance of individual metabolites is in Figure S1. One-way ANOVA (*n* = 4 biological replicates per condition). * p ≤ 0.05; ** p ≤ 0.01; *** p ≤ 0.001; **** p ≤ 0.0001

**Table 1.** Mucin Glycan, Protein, and Iron Content.

| | Glycans<br>(mg/mL) | Protein<br>(mg/mL) | Glycans/<br>Protein | Iron<br>(μM)<sup>#</sup> |
| --- | --- | --- | --- | --- |
| PGM | 2.82 | 0.96 | 3.0 | 33.6 |
| dPGM | 1.51 | 0.54 | 2.8 | 14.9 |
| cPGM | 2.42 | 0.53 | 4.6 | 2.6 |
| dcPGM | 1.33 | 0.30 | 4.4 | 1.1 |
| BSM | 2.17 | 0.58 | 3.7 | 14.6 |
| dBSM | 0.25 | 0.23 | 1.1 | 3.7 |

d: dialyzed; c: clarified; <sup>#</sup>in 5 mg/mL material

Secondary metabolite profiling of the cultures was performed using liquid chromatography tandem mass spectrometry and individual metabolites were grouped into structurally related molecular families (Figure 1B, Figure S1).(11) At the molecular family level, PAO1 in PGM-SCFM2 produced 2.5-fold higher levels of phenazines (PHZs), 2-fold lower levels of rhamnolipids (RLs), and the levels of the siderophores pyochelin (PCH) and pyoverdine (PVD) were measured at the limit of detection, while in BSM-SCFM2 PAO1 produced 1.2-fold lower amounts of RLs, 1.4-fold higher amounts of PCH, and 7.7-fold lower amounts of PVD compared to None-SCFM2 cultures (Figure 1B). No apparent difference in abundance of the alkyl quinolone (AQ) molecular family was observed between the three conditions.

Broadly, this pattern of secondary metabolite production indicates that the primary response of PAO1 to BSM and PGM in SCFM2 is driven by their different iron concentrations. In *P. aeruginosa*, secondary metabolite production is regulated, in part, by iron environment concentration through the ferric uptake regulator (Fur).(31, 39, 40) In iron deplete conditions, *P. aeruginosa* produces the siderophores PCH and PVD scavenge ferric (Fe^3+^) iron, forming soluble complexes that are taken up into the cell. *P. aeruginosa* also upregulates RL production via Fur to support swarming motility, biofilm formation, enhance microbial competition, and lyse cells.(41, 42) In iron replete conditions, iron-bound Fur represses the biosynthesis of these molecular families. In aerobic environments, Fe^2+^ reacts with environmental oxygen to form insoluble Fe^3+^. In response, *P. aeruginosa* increases production of PHZs to reduce insoluble Fe^3+^ to soluble Fe^2+^ for uptake via the ferrous iron transport (Feo) system.(43)

The secondary metabolite profiling data confirmed our previous results that addition of PGM to SCFM2 creates an iron-replete environment, marked by reduced production of RLs and siderophores, likely through repression of their biosynthetic pathways by Fur, and enhanced production of PHZs to reduce insoluble iron.(11) This data also revealed that addition of BSM to SCFM2 creates an environment containing moderate concentrations of iron compared to the low iron None-SCFM2, with increased levels of PCH and decreased levels of PVDs indicative of siderophore switching by PAO1.(44) *P. aeruginosa* uses siderophore switching to modulate PCH and PVD levels in response to alterations in iron availability. Due to its higher affinity for iron, PVD is considered the primary siderophore of *P. aeruginosa* and is only produced under extremely low iron conditions, such as those modeled by None-SCFM2.(45, 46) In PAO1, PVD biosynthesis is decreased in response to the presence of increasing iron concentration, with production of its secondary siderophore PCH maximized in environmental conditions containing ∼10 µM iron.(44) This nuanced approach to iron acquisition by *P. aeruginosa* enables it to survive in a myriad of environments.

### Generation of low-iron commercial mucin

The concentration of iron in commercial mucins has implications in appropriately modeling the nutritional environment of the CF airway. The 5 mg/mL mucin added to SCFM2 represents the lowest concentration of mucin added to ASM, with some formulations supplementing the media with 20 mg/mL to better mimic the mucin concentrations measured from CF populations of advanced age and/or worsened airway disease.(11, 12) However, the mean iron concentration measured from the CF airway ranges from ∼4 to 35 μM, with higher levels associated with decreased lung function and increased *P. aeruginosa* burden.(47, 48) In the models of ASM with 20 mg/mL of PGM added to the medium, the iron concentration is over 100 µM, well above the physiological concentrations of the CF airway. The intrinsically linked concentration of iron with the amount of mucin added to the medium complicates the ability to differentiate the effect of iron concentration and mucin structure on microbial physiology and community interactions under physiologically relevant conditions *in vitro*.

Multiple approaches have been developed to purify mucin from commercial preparations, primarily for biophysical studies.(13, 17, 49–53) As our goal was to simply remove iron from the commercial mucins, we applied ultracentrifugation and dialysis; two broadly accessible methods for semi-purification of proteins. The iron concentration of the crude mucin and subsequent preparations was measured by inductively coupled plasma mass spectrometry (ICP-MS). Whereas we used ICP-MS to quantify the iron concentration of the mucin preparation, a commercial kit can be used with appropriate dilution of the mucin preparations to iron concentrations within the linear range of the assay. Mucin purity was measured as the ratio of glycan concentration to total protein.(54) Unfortunately, ELISA-based quantification of MUC5AC and MUC5B from the crude PGM and BSM was unsuccessful, likely due to loss of the C-termini during commercial processing.(13, 55) During this analysis, we observed that the iron concentration and mucin purity of crude commercial mucins varied significantly between types (e.g., PGM vs BSM), manufacturers (e.g., Sigma vs Lee), lots, and within bottles due to variations in commercial processing and the presence of microbial contaminants. Therefore, it is necessary to quantify both the iron concentration and mucin purity of semi-purified mucin prior to use.

To reduce the iron concentration of PGM, we clarified soluble mucin and removed unbound iron by dialysis.(17) This method of semi-purification of PGM reproducibly resulted in low iron mucin preparations. First, cellular debris was removed from PGM using ultracentrifugation. Clarified PGM (cPGM) contained 2.6 µM iron with a 1.5-fold increase in mucin purity. Subsequent dialysis of cPGM (dcPGM) against a 10 kD molecular weight cutoff (MWCO) filter further reduced the iron concentration to 1.1 µM. However, dialysis led to a 1.8-fold decrease in both glycan and protein concentration. Considering the molecular weight of monomeric and polymeric mucin is ∼650 kD and 1-4 MDa, the reduction in glycan and protein concentration suggested that roughly half of the crude PGM preparation was comprised of small, glycosylated peptides/proteins.(56, 57) Indeed, dialysis of PGM (dPGM) led to a similar 1.8-fold reduction in glycan and protein levels. Likewise, dialysis of BSM (dBSM) against a 10 kD MWCO filter led to an 8.7-fold reduction in the glycan concentration and a 2.5-fold loss of total protein, which suggested that BSM was primarily comprised of mucin degradation products. Further purification of BSM was not pursued due to the cost of the material.

As ultracentrifugation was sufficient to reduce the concentration of iron in PGM to below physiologically relevant levels while improving mucin purity, we did not pursue full purification of the polymeric mucin. As a result, some soluble non-mucin components, including proteins and DNA, are likely in our preparation. If desired, these non-mucin components can be removed by published methods for mucin purification.(17) Based upon the concomitant decrease in glycan and protein concentration, we surmised that the material lost during dialysis represents small, glycosylated peptides and proteins from degradation of the mucins during commercial processing. How these mucin degradation products influence the experimental results of microbiology experiments is unknown. However, in the context of the CF airway, inclusion of the glycopeptides in *in vitro* model systems may better represent the highly proteolytic *in vivo* environment.(58) Therefore, we used cPGM as the mucin additive to SCFM2 for all additional experiments.

### Iron and mucin structure modulate PAO1 secondary metabolite production in SCFM2

To determine whether iron concentration was indeed the driving factor of the enhance growth and secondary metabolite profile observed for PAO1 in PGM-SCFM2, comparative metabolomics of cultures in PGM-SCFM2, cPGM-SCFM2, and cPGM-SCFM2 complemented with FeSO_4_ (cPGM+Fe-SCFM2) was performed. PGM- and cPGM-SCFM2 were formulated to contain equivalent concentrations of soluble mucin based on mucin purity. Additionally, PGM- and cPGM+Fe-SCFM2 were formulated to contain equivalent total iron concentrations. Enumeration of viable cells after 48 hr of static incubation revealed that PGM-SCFM2 provided a slightly more advantageous growth environment for PAO1, resulting in a ∼4-fold higher recovery of CFU/mL than the other two culture conditions (Figure 2A). The result indicates that nutrients other than soluble mucin and iron are contributing to enhanced growth of PAO1 cultures supplemented with crude PGM.

**Figure 2.**
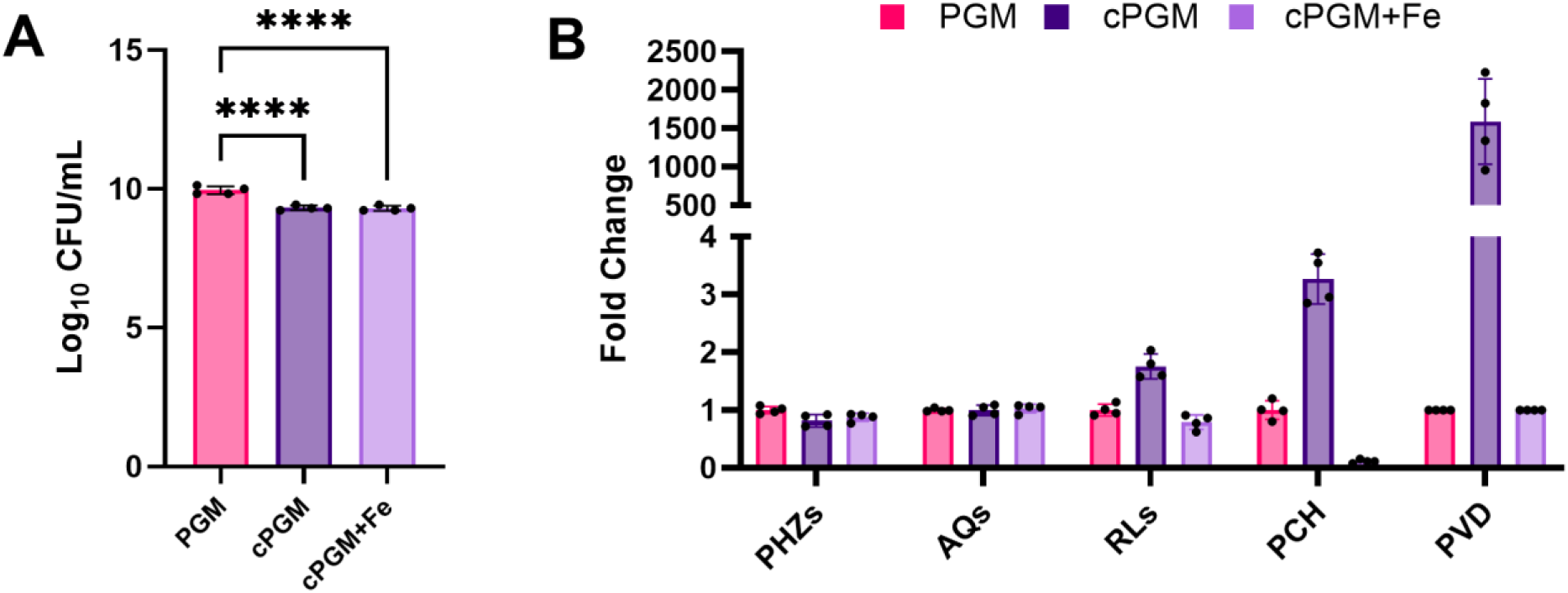
(A) Viable cell counts (CFU/mL) from *P. aeruginosa* PAO1 48 hr static cultures in PGM-SCFM2, cPGM-SCFM2, and cPGM-SCFM2+Fe. PGM-SCFM2 and cPGM-SCFM2 contain equal concentrations of mucin. PGM-SCFM2 and cPGM-SCFM2+Fe contain equal concentrations of iron. (B) Fold change secondary metabolite levels produced by PAO1 in cPGM- and cPGM+Fe-SCFM2 compared to PGM-SCFM2. PGM: porcine gastric mucin; cPGM: clarified PGM; cPGM+Fe: clarified PGM supplemented with iron; PHZs: phenazines; AQs: alkyl quinolones; RLs: rhamnolipids; PCH: pyochelin; PVDs: pyoverdines. Corresponding abundance of individual metabolites is in Figure S2. One-way ANOVA (*n* = 4 biological replicates per condition). * p ≤ 0.05; ** p ≤ 0.01; *** p ≤ 0.001; **** p ≤ 0.0001

Secondary metabolite profiling revealed that PAO1 was canonically responsive to iron concentration under these culture conditions (Figure 2B, Figure S2).(11) Indicative of its ∼6 µM total iron concentration, PAO1 produced 1.25-lower levels of PHZs, 1.75-fold higher levels of RLs, 3.2-fold higher levels of PCH, and ∼1500-fold higher levels of PVD in cPGM-SCFM2 compared to PGM-SCFM2 cultures. Addition of Fe^2+^ to cPGM decreased the levels of RLs, PCH, and PVD produced by PAO1 to those at or below the amounts measured from the PGM-SCFM2 cultures. Although PGM- and cPGM+Fe-SCFM2 contain equivalent total iron concentration, PAO1 produced lower levels of PCH in cPGM+Fe-SCFM2, likely due to higher levels of unbound Fe^2+^.

Due to the excess of Fe^2+^ in cPGM+Fe-SCFM2, we expected that PAO1 would produce higher quantities of PHZs to reduce insoluble Fe^3+^ in this condition than in the low iron conditions of cPGM-SCFM2.(43) However, total PHZ levels were equivalent (Figure 2B). Quantitation of the individual PHZs pyocyanin (PYO), phenazine-1-carboxamide (PCN), and phenazine-1-carboxylic acid (PCA) revealed that while PYO levels were lower in cPGM+Fe-SCFM2 cultures compared to cPGM-SCFM2 cultures, increased PCA levels in response to higher levels of Fe^3+^ compensated for this decrease leading to similar levels of total PHZs (Figure S2).(43) The slight alterations in the levels of individual PHZs produced by PAO1 under these culture conditions did not align with the trend of markedly higher (2 to 3-fold) levels of PHZs produced in PGM-SCFM2 compared to both None- and BSM-SCFM2 (Figure 1B, Figure S1). Rather, the level of PHZs remained elevated in all media containing any derivative of PGM regardless of the total iron concentration, indicating the mucin structure of PGM induces higher PHZ production by PAO1.

Despite inclusion of both BSM and PGM in media aiming to model mucosal environments, they are comprised of different mucins. BSM is comprised of MUC5B, whereas PGM is predominantly MUC5AC.(11, 17–21, 24) Consequently, PGM and BSM exhibit several structural differences, including different domain organization, variation in monosaccharide incorporation frequency, and terminal sulfation, fucosylation, and sialylation of the O-glycans.(59) One of the structural differences between MUC5AC (PGM) and MUC5B (BSM) is the level of *N-*acetylglucosamine (GlcNAc) incorporation. GlcNAc constitutes ∼39% of the monosaccharides present in MUC5AB, but only ∼19% of the MUC5B monosaccharides. Microarray analysis of *P. aeruginosa* cultured in sputum has revealed an increase in the transcription of genes involved in GlcNAc catabolism, indicating it is likely an important carbon source in the infection environment of CF lungs.(59) Reflecting this significance, GlcNAc is included in SCFM2 as a nutrient source.(33) *In vitro*, GlcNAc has been shown to increase the production of PHZs by *P. aeruginosa*. (60) Therefore, the elevated PHZs levels in PGM containing media could be caused by the higher incorporation rate of the GlcNAc into MUC5AC. Importantly, the higher levels of PHZs produced by P. aeruginosa in culture media containing PGM will impact the results of antibiotic sensitivity testing due to the role of PHZs in promoting tolerance to clinically relevant antibiotics.(61)

### Iron concentration modulates competition between *P. aeruginosa* and *S. aureus* within a CF SymCom in cPGM-SCFM2

Although *P. aeruginosa* is one of the primary pathogens colonizing the airways of people with CF, the CF airways are host to a diverse microbiota.(62, 63) This dynamic microbial community comprised of obligate and facultative anaerobes from the oropharynx and canonical CF pathogens, such as *P. aeruginosa* and *S. aureus*, form stable, localized communities due, in part, to metabolic cross-feeding of glycans and amino acids from mucin-degraders, including *Prevotella* spp.(5, 8, 64) These mucin-dependent interactions of the CF airway microbiota influence infection outcomes by promoting the establishment of chronic infection by opportunistic pathogens and alter sensitivity to antimicrobial therapy.(65)

Recently, a genetically tractable, four-member model CF synthetic community created from microbial profiling of CF airway samples was created. This community was comprised of *P. aeruginosa*, *S. aureus*, *Streptococcus sanguinis*, and *Prevotella melaninogenica* and stably modeled in PGM-SCFM2 under anoxic conditions.(29) *P. aeruginosa* secondary metabolites have been detected in CF airway samples and biosynthesis of those metabolites requires oxygen, suggesting that some *P. aeruginosa* reside in aerobic or microaerophilic pockets within the CF airways. (66–71) Therefore, we were curious whether the composition of the 4-member CF SynCom could be recapitulated under atmospheric conditions and if the iron content of PGM modulated community structure by decreasing competition for this nutrient.

The mixed culture of *P. aeruginosa* PA14, *S. aureus* USA300, *S. sanguinis* SK36, and *P. melaninogenica* ATCC25845 and associated monocultures were grown in PGM-SCFM2 and cPGM-SCFM2 under static, aerobic conditions for 48 hr. Aligning well with reported composition, viable cell counts were recovered for all four species from the mixed SynCom cultures, with enhanced growth of *S. sanguinis* and *P. melaninogenica* compared to monocultures (Figure 3A).(29) Under the low iron conditions of cPGM-SCFM2, *S. aureus* was not recovered from the SynCom cultures. Since *S. aureus* grew robustly in cPGM-SCFM2 monocultures (Figure 3B), it was evident that the decreased recovery of *S. aureus* from the community was not due to a growth defect in the medium, but interspecies competition. Since the antagonistic interactions between *P. aeruginosa* and *S. aureus* are well characterized, we hypothesized that *P. aeruginosa* anti-staphylococcal small molecule virulence factors would be produced in higher quantity in cPGM-SCFM2.

**Figure 3.**
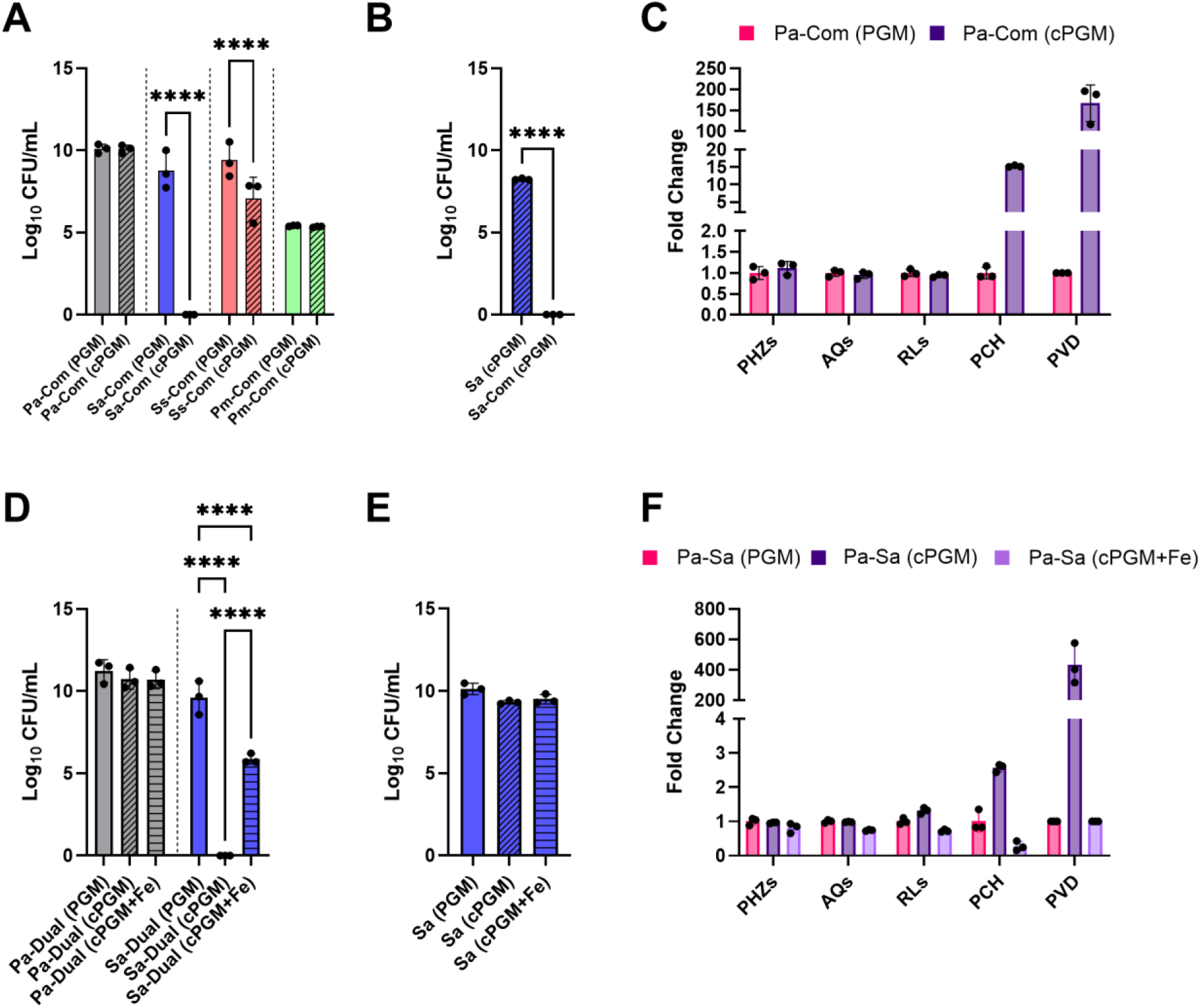
(A) Viable cell counts (CFU/mL) of *Pseudomonas aeruginosa* PA14 (Pa), *Staphylococcus aureus* USA300 (Sa), *Streptococcus sanguinis* SK36 (Ss), *Prevotella melaninogenica* ATCC25845 (Pm) from 48 hr static CF SynCom (Com) cultures in PGM-SCFM2 (PGM) and cPGM-SCFM2 (cPGM). Viable cell counts from all mono- and mixed cultures are in Figure S3. (B) Viable cell counts (CFU/mL) of *S. aureus* USA300 (Sa) from 48 hr static monocultures and CF SymCom (Com) in cPGM-SCFM2 (cPGM). (C) Fold change secondary metabolite levels produced by *P. aeruginosa* PA14 in 48 hr static CF SynCom (Com) cultures in cPGM-SCFM2 (cPGM) compared to PGM-SCFM2 (PGM). Corresponding abundance of individual metabolites is in Figures S4. (D) Viable cell counts (CFU/mL) of *P. aeruginosa* PA14 (Pa) and *S. aureus* USA300 (Sa) from 48 hr static co-cultures (Dual) in PGM-SCFM2 (PGM), cPGM-SCFM2 (cPGM), and cPGM+Fe-SCFM2. Viable cell counts from all mono- and co-cultures are in Figure S5. (E) Viable cell counts (CFU/mL) of *S. aureus* USA300 (Sa) from 48 hr static monocultures in PGM-SCFM2 (PGM), cPGM-SCFM2 (cPGM), and cPGM+Fe-SCFM2. (F) Fold change secondary metabolite levels produced by *P. aeruginosa* PA14 in 48 hr static co-culture with *S. aureus* USA300 in cPGM-SCFM2 (cPGM) and cPGM+Fe-SCFM2 (cPGM+Fe) compared to PGM-SCFM2 (PGM). PGM: porcine gastric mucin; cPGM: clarified PGM; cPGM+Fe: clarified PGM supplemented with iron; PHZs: phenazines; AQs: alkyl quinolones; RLs: rhamnolipids; PCH: pyochelin; PVDs: pyoverdines. Corresponding abundance of individual metabolites is in Figures S6. Paired t test (A and B) or one-way ANOVA (C or D) (*n* = 3 biological replicates per condition). * p ≤ 0.05; ** p ≤ 0.01; *** p ≤ 0.001; **** p ≤ 0.0001

Several secondary metabolites have been shown to contribute to the ability of *P. aeruginosa* to outcompete *S. aureus*.(72) These mechanisms include inhibition of the electron transport chain by the AQ HQNO, dispersal of biofilm by RLs, iron sequestration by the siderophores PCH and PVD, and induction of intracellular reactive oxygen species (ROS) by PYO and PCH.(73–77) Comparative metabolomics analysis of the CF SynCom cultured in PGM- and cPGM-SCFM2 revealed that *S. aureus* was outcompeted by *P. aeruginosa* through iron restriction by the siderophores PCH and PVD (Figure 3C, Figure S4). These results indicated that *P. aeruginosa* siderophore production in response to limited environmental iron was responsible for inhibition of *S. aureus* within the CF SynCom cultured in cPGM-SCFM2.

To confirm this observation, *P. aeruginosa* PA14 and *S. aureus* USA300 were co-cultured in PGM-, cPGM-, and cPGM+Fe-SCFM2. As in the CF SynCom, *S. aureus* was not recovered from the cPGM-SCFM2 co-cultures (Figure 3D, Figure S5). The lower recovery of *S. aureus* from cPGM-SCFM2 was not due to reduced viability (Figure 3E). Secondary metabolite profiling of the co-cultures confirmed higher levels of PCH and PVD in cPGM-SCFM2 (Figure 3F, Figure S6). Reflective of the low iron concentration of cPGM-SCFM2, RL levels were also slightly higher, but the amounts of PHZs and AQs were equivalent between the two conditions. Although addition of Fe^2+^ in cPGM-SCFM2 led to levels of PCH, PVD, and RLs at or below those measured from PGM-SCFM2 cultures, recovery of *S. aureus* from cPGM+Fe-SCFM2 co-cultures was ∼4 log lower, despite equivalent concentrations of total iron in PGM and cPGM+Fe (Figure 3D). This growth defect was not observed in monoculture (Figure 3E).

Together, our data illustrates that commercial mucins contain undefined sources of iron that impact the physiology of individual microorganisms in both monoculture and mixed communities, confounding the interpretation of experimental results within media aiming to model mucosal environments. This work illustrates that the impact of commercial mucins on *P. aeruginosa* in both monocultures and within the CF SymCom is multifactorial. The iron content of commercial mucins influences the levels of secondary metabolites produced by *P. aeruginosa*, primarily reduction of its siderophores PCH and PVD and to a lesser extent RLs. The structural differences between BSM and PGM contribute to differential production of PHZs, which may influence the outcomes of both interspecies interactions as antimicrobial susceptibility testing. Additionally, undefined components of crude PGM that are removed during clarification enhance the co-existence of *S. aureus* with *P. aeruginosa*. Although our experiments evaluated the role of iron in SCFM2 as a model system of the CF airway, media formulations aiming to replicate the gut mucosal environment, including complex intestinal medium (CIM) and gut microbiota medium (GMM), also contain undefined sources of iron such as PGM (type II) and meat extract, respectively, indicating that the influence of undefined iron in media components used in *in vitro* modeling of mucosal microbiota is wide-spread, reducing competition for iron, and potentially influencing experimental interpretation of microbial interactions.(10, 78) The ability to control the iron concentration in mucosal media will enable researchers to evaluate the iron content of different disease states on microbial community composition and pathogen expansion as well as more accurately model the *in vivo* environment by mirroring nutritional immunity by incorporating host-derived iron binding proteins under physiologically relevant conditions.

## MATERIALS AND METHODS

### Mucin preparation

Commercial porcine gastric mucin type III (PGM, Millipore Sigma) or bovine submaxillary mucin (BSM, Millipore Sigma) was suspended in 1X 3-morpholinopropane-1-sulfonic acid (MOPS) buffer (pH 7.0) at 5% (w/v). Dialysis of the crude mucins was performed using 10 kD molecular weight cutoff cellulose membrane cassettes (Thermo Scientific) against 1X MOPS. Clarification was performed by ultracentrifugation as previously described with minor modifications.(17) PGM suspended in 10 mM MOPS buffer was centrifuged at 8,300 x g for 30 minutes at 4 °C (Thermo Sorvall ST 40R). Subsequently, the supernatant was transferred to a clean centrifuge tube and centrifuged again at 15,000 x g for 45 minutes at 4 °C (Sorvall RC5C). The mucin preparations were sterilized using a liquid autoclave cycle (<20 min sterilization time), stored at 4 °C, checked for sterility, and used within 1 month of preparation.

### Mucin quantification

Mucin purity was quantified as the ratio of glycan concentration to the total protein concentration. Glycans were quantified using the periodic acid – Schiff (PAS) stain assay as detailed by Kilcoyne, *et al*. (54) Briefly, 25 µl of each sample was added to wells of a 96 well plate and 120 µl periodic acid solution (0.06% periodic acid in 7% acetic acid) was added and mixed by pipetting. The plate was covered with a plastic seal and incubated at 37 °C for 90 min. The plate was brought to room temperature and 100 µl Schiff’s reagent (Millipore Sigma) was added and mixed by pipetting. The plate was covered with a plastic seal, was shaken 5 min, then left at room temperature for 40 min. The seal was removed, and the plate was read at 550 nm using a Synergy HT plate reader (BioTek). Glycan concentration was quantified using an *N*-acetylgalactosamine (MP Biomedicals) standard curve collected on the same plate. Total protein concentration was quantified by the Pierce BCA Protein Assay Kit (Thermo Scientific) following manufacturer instructions.

### Iron quantitation

Suspensions of processed and unprocessed PGM and BSM in 1X MOPS were diluted 1:1 with trace metal grade nitric acid, digested overnight, and diluted 10-fold. Indium-3, 5 ppb, was added to each sample as an internal standard. Metal concentration was measured by inductively coupled plasma mass spectrometry (ICP-MS) analysis on a PerkinElmer NexION 2000B. Samples were analyzed in technical triplicate with a blank wash run between each sample. Data were collected using the sample acquisition module within Syngistix software (version 2.3) and analyzed using Microsoft Excel.

### Microbial culture

Bacterial strains: *P. aeruginosa* PAO1 (MPAO1) - University of Washington; *P. aeruginosa* PA14 - George O’Toole (Dartmouth Geisel School of Medicine); *S. aureus* USA300 TCH1516 - Victor Nizet (University of California, San Diego); *S. sanguinis* SK36 - Jens Kreth (Oregon Health & Science University); *P. melaninogenica* ATCC25845 (ATCC). Synthetic Cystic Fibrosis Medium 2 (SCFM2) was prepared as previously described with the modifications detailed below. (33)

#### *P. aeruginosa* PAO1 in SCFM2 complemented with PGM or BSM

*P. aeruginosa* strain PAO1 was inoculated from a streak plate into 5 mL LB and incubated overnight at 37 °C, shaking at 220 RPM. One milliliter of SCFM2 without mucin (None-SCFM2), SCFM2 complemented with 5 mg/mL PGM (PGM-SCFM2), and SCFM2 complemented with 5 mg/mL BSM (BSM-SCFM2) were inoculated with ∼1×10^6^ CFU/mL PAO1 in a polystyrene 48 well plate. The plate was covered with its lid and incubated statically, under ambient oxygen conditions, at 37 °C for 48 hr. Samples were mechanically disrupted. An aliquot was retained for metabolomics analysis. Enumeration of viable cells (CFU/mL) was performed by serial dilution on LB agar (Millipore Sigma) incubated at 37 °C.

#### *P. aeruginosa* PAO1 in SCFM2 complemented with PGM, cPGM, or cPGM+Fe

*P. aeruginosa* PAO1 was inoculated from a streak plate into 5 mL LB and incubated overnight at 37 °C, shaking at 220 RPM. One milliliter of SCFM2 complemented with 5 mg/mL PGM (PGM-SCFM2), SCFM2 complemented with 3.26 mg/mL cPGM (cPGM-SCFM2), and SCFM2 complemented with 3.26 mg/mL cPGM and 31 µM FeSO_4_ (cPGM+Fe-SCFM2) were inoculated with ∼1×10^6^ CFU/mL PAO1 in a polystyrene 48 well plate. The media PGM- and cPGM-SCFM2 contained equivalent amounts of mucin as measured by the glycan to protein ratio. The media PGM- and cPGM+Fe-SCFM2 contain equimolar concentrations of iron. The plate was covered with its lid and incubated statically, under ambient oxygen conditions, at 37 °C for 48 hr. Samples were mechanically disrupted. An aliquot was retained for metabolomics analysis. Enumeration of viable cells (CFU/mL) was performed by serial dilution on Pseudomonas isolation agar (PIA, BD Difco) incubated at 37 °C.

#### CF SynCom in SCFM2 complemented with PGM or cPGM

The four member CF synthetic community was cultured as previously described, with slight modifications.(29) *P. aeruginosa* PA14 and *S. aureus* USA300 were inoculated from a streak plate into 5 mL tryptic soy broth (TSB, BD Difco) and incubated overnight at 37 °C, shaking at 220 RPM. *S. sanguinis* SK36 was inoculated from a streak plate into 5 mL of Todd-Hewitt broth (BD Difco) supplemented with 0.5% yeast extract (BD Difco) and incubated statically overnight at 37 °C in a 5% CO_2_ atmosphere. *P. melaninogenica* ATCC25845 was inoculated into TSB supplemented with 0.5% yeast extract, 5 µg/mL hemin (Millipore sigma), 2.85 mM L-cysteine (Acros Organics), and 1 µg/mL menadione (United States Pharmacopeia Convention) and incubated statically overnight at 37 °C in an anoxic atmosphere (BD GasPak). *S. sanguinis* and *P. melaninogenica* cultures were centrifuged and the cell pellets were washed once with sterile 1X PBS. *P. aeruginosa* and *S. aureus* cultures were centrifuged, and the cell pellets were washed twice with sterile 1X PBS. All four species were diluted to an OD_600_ of 0.2 in None-SCFM2. For mixed culture, 80 µL of PGM-SCFM2 or cPGM-SCFM2 were inoculated with 5 µL of each diluted culture in a sterile 96 well polypropylene plate (Nunc). For monoculture, 95 µL of PGM-SCFM2 and cPGM-SCFM2 were inoculated with 5 µL of individual diluted cultures. The plate was incubated statically at 37 °C under ambient atmosphere for 48 hr. Samples were mechanically disrupted. An aliquot was retained for metabolomics analysis. Enumeration of viable cells (CFU/mL) was performed by serial dilution as follows: (1) *P. aeruginosa* - from PIA after overnight incubation at 37 °C under ambient atmosphere; (2) *S. aureus* - from mannitol salt agar (BD Difco) after overnight incubation at 37 °C under ambient atmosphere; (3) *S. sanguinis* - from TSB supplemented with 0.5% yeast extract, 1.5% agar (BD Difco), 5% sheep’s blood (Remel, Thermo Scientific), 10 µg/mL oxolinic acid (Acros Organics), 10 µg/mL polymixin B (Alfa Aesar) after overnight incubation at 37 °C under a 5% CO_2_ atmosphere; (4) *P. melaninogenica* - from TSB supplemented with 0.5% yeast extract, 1.5% agar (BD Difco), 5% sheep’s blood, 5 µg/mL hemin, 2.85 mM L-cysteine, 1 µg/mL menadione, 5 µg/mL vancomycin (Alfa Aesar), and 100 µg/mL kanamycin (Fisher Scientific) after overnight incubation at 37 °C under an anoxic atmosphere.

#### *P. aeruginosa* PA14 and *S. aureus* USA300 in SCFM2 complemented with PGM, cPGM, or cPGM+Fe

*P. aeruginosa* strain PA14 and *S. aureus* strain USA300 were inoculated from a streak plate into 5 mL TSB and incubated overnight at 37 °C, shaking at 220 RPM. *P. aeruginosa* and *S. aureus* cultures were centrifuged, and the cell pellets were washed twice with sterile 1X PBS then diluted to an OD_600_ of 0.2 in None-SCFM2. For co-culture, 90 µL of PGM-SCFM2, cPGM-SCFM2, or cPGM-SCFM2+Fe was inoculated with 5 µL of each diluted culture in a sterile 96 well polypropylene plate (Nunc). For monoculture, 95 µL of PGM-SCFM2, cPGM-SCFM2, or cPGM-SCFM2+Fe was inoculated with 5 µL of individual diluted cultures. The plate was incubated statically at 37 °C under ambient atmosphere for 48 hr. Samples were mechanically disrupted. An aliquot was retained for metabolomics analysis. Enumeration of viable cells (CFU/mL) was performed by serial dilution as follows: (1) *P. aeruginosa* - from PIA after overnight incubation at 37 °C under ambient atmosphere; (2) *S. aureus* - from mannitol salt agar after overnight incubation at 37 °C under ambient atmosphere.

### Metabolomics sample preparation

#### P. aeruginosa PAO1 samples

Each sample was chemically disrupted with an equal volume of 1:1 solution of ethyl acetate (EtOAc, VWR HiPerSolv Chromanorm) and methanol (MeOH, Fisher Scientific Optima LC/MS grade).

#### Mixed and co-culture samples

Each sample was chemically disrupted with an equal volume of methanol containing 10 µM nalidixic acid (Thermo Scientific). All samples were dried and stored at −20°C until use. After thawing, samples were resuspended in 50% MeOH, diluted 10-fold in 50% MeOH containing 1 µM glycocholic acid (Calbiochem, 100.1% pure), and centrifuged for 10 min at 4000 RPM (Thermo Sorvall ST 40R) to remove non-soluble particulates prior to injection.

### LC-MS/MS Data Acquisition

Mass spectrometry data acquisition was performed using a Bruker Daltonics Maxis II HD qTOF mass spectrometer equipped with a standard electrospray ionization (ESI) source as previously described.(11) The mass spectrometer was tuned by infusion of Tuning Mix ESI-TOF (Agilent Technologies) at a 3 µL/min flow rate. For accurate mass measurements, a wick saturated with Hexakis (1H,1H,2H-difluoroethoxy) phosphazene ions (Apollo Scientific, *m/z* 622.1978) located within the source was used as a lock mass internal calibrant. Samples were introduced by an Agilent 1290 UPLC using a 10 µL injection volume.

Extracts were separated using a Phenomenex Kinetex 2.6 µm C18 column (2.1 mm x 50 mm) using a 9 minute, linear water-ACN gradient (from 98:2 to 2:98 water:ACN) containing 0.1% FA at a flow rate of 0.5 mL/min. The mass spectrometer was operated in data dependent positive ion mode, automatically switching between full scan MS and MS/MS acquisitions. Full scan MS spectra (*m/z* 50 - 1500) were acquired in the TOF and the top five most intense ions in a particular scan were fragmented via collision induced dissociation (CID) using the stepping function in the collision cell. LC-MS/MS data for PA mix, a mixture of available commercial standards of *P. aeruginosa* secondary metabolites, were acquired under identical conditions. Bruker Daltonics CompassXport was used to apply lock mass calibration and convert the LC-MS/MS data from proprietary to open-source format.

### *P. aeruginosa* metabolite quantitation and annotation

MZmine (version 3.9.0) was used to perform feature finding on the. mzML files as previously described.(11, 79) Features were normalized by row sum then filtered for *P. aeruginosa* secondary metabolites using exact mass and retention time. Annotation of the phenazines, alkyl quinolones, and rhamnolipids was manually confirmed by comparing the experimental data (exact mass, MS/MS, and retention time) with data corresponding to commercial standards (level 1 annotation) using GNPS Dashboard and Metabolomics Spectrum Resolver.(80, 81) Annotation of pyochelin and pyoverdine was manually confirmed by comparing experimental data (exact mass, MS/MS) to spectra deposited into the GNPS libraries (level 2 annotation) and reported structures.(82–85)

### Statistical analysis

Statistical comparison of CFU and secondary metabolite levels between sample types was conducted in GraphPad Prism (version 9.3.1) as described in the figure legends. For all analyses, p < 0.05 were considered statistically significant.

## Data availability

All mass spectrometry data are available at MassIVE: PAO1 in None-, PGM-, and BSM-SCFM2 (MSV000095750); PAO1 in PGM-, cPGM, cPGM+Fe-SCFM2 (MSV000095752); 4-member SynCom in PGM- and cPGM-SCFM2 (MSV000095754); PAO1-USA300 in PGM-, cPGM-, and cPGM+Fe-SCFM2 (MSV000095755). For review, password is maldi3480

## ACKNOWLEDGEMENTS

This work was supported by NIGMS grant R35 GM128690 and the ALSAM Foundation to V.V.P. We are grateful to Justin Jens and Matilda Fiebig for performing initial experiments.

## AUTHOR CONTRIBUTIONS

EG, RLN, and VVP designed and performed the research, analyzed the data, and wrote the manuscript.

## COMPETING INTERESTS

The authors declare that there are no competing interests.

